# Extracellular Vesicles from Senescent Tumor Cells Are Necessary and Sufficient to Drive Paracrine Senescence

**DOI:** 10.64898/2026.03.25.713920

**Authors:** Valentin Estevez-Souto, Alex Miralles-Dominguez, Pablo Pedrosa, Patricia Lado-Fernandez, Miguel A. Prados, Alejandro Failde-Fiestras, Raquel Paredes-Paredes, Jorge Ruz-Ortega, Maria J. Alonso, Martina Migliavacca, Ester Polo, Rebeca Alvarez-Velez, Enrique Vazquez de Luis, Ana Dopazo, Gabriela N. Condezo, Carmen San Martin, Miguel Gonzalez-Barcia, Pilar Ximénez-Embún, Javier Muñoz, Manuel Collado, Sabela Da Silva-Alvarez

## Abstract

Cellular senescence exerts powerful non-cell autonomous effects through the senescencelzlassociated secretory phenotype (SASP). This SASP comprises soluble factors and extracellular vesicles (EVs). Although soluble SASP components can induce senescence in neigbouring cells, the specific contribution of EVs to paracrine senescence is poorly defined. Here, we show that EVs released by senescent tumor cells are necessary and sufficient to propagate senescence. Conditioned media from bleomycinlzlinduced senescent A549 cells triggered a permanent growth arrest with morphological changes and upregulation of senescence markers in recipient tumor cells. Pharmacological inhibition of EV biogenesis using GW4869 or genetic downregulation of the EV secretion mediator RAB27A markedly attenuates paracrine senescence without affecting soluble SASP factor secretion or the senescent state of producer cells. Proteomic characterization reveals that senescent EVs exhibit a distinct molecular signature enriched for extracellular components and processes related to wound healing and hemostasis. Importantly, purified senescent EVs, devoid of soluble SASP factors, fully recapitulated paracrine senescence induction. These findings identify senescent EVs as key autonomous SASP effectors and highlight vesicular pathways as potential therapeutic targets in cancer and therapylzlinduced senescence.

## INTRODUCTION

Cellular senescence, a state of stable cell cycle arrest, represents a fundamental tumor suppressive mechanism that prevents the proliferation of damaged or oncogene-activated cells (Collado et al., 2007; Serrano, 2011). Paradoxically, senescent cells can persist in tissues and secrete a complex repertoire of factors collectively termed as the senescence-associated secretory phenotype (SASP), which can exert pro-tumorigenic effects through the remodeling of the microenvironment (Acosta et al., 2013; Coppé et al., 2010). The SASP comprises a complex mixture of soluble factors such as cytokines, chemokines, growth factors, and extracellular matrix remodelers with functional consequences that vary depending on the context and that might result in beneficial or detrimental effects. Recently, it has become apparent that apart from this soluble SASP there is also a vesicular SASP, formed mainly by extracellular vesicles (EVs). Potentially, both the soluble and the vesicular fraction can contribute to the heterogeneous functional outcomes of senescence in cancer progression promoted by the SASP (Borghesan et al., 2019).

While the soluble fraction of the SASP has been extensively characterized for its capacity to induce senescence in neighboring cells, a phenomenon termed paracrine senescence, the specific contribution of EVs to this process remains incompletely understood. EVs serve as critical mediators of intercellular communication by transferring bioactive cargo, including proteins, lipids, and nucleic acids, between cells (Raposo & Stoorvogel, 2013; Welsh et al., 2024). Recent studies have revealed that senescent cells release increased quantities of EVs, which have been implicated in diverse biological processes ranging from immune modulation to tissue repair (Estévez-Souto et al., 2023; Takasugi, 2018). However, whether senescent EVs possess autonomous paracrine senescence-inducing activity, distinct from soluble SASP components, has not been rigorously tested.

The functional dissection of SASP components carries significant therapeutic implications. Pharmacological strategies targeting senescent cells using compounds that induce specific cytotoxicity, known as senolytics, or that dampen down their secretory output, known as senomorphics, are actively being developed for cancer and age-related pathologies (Hickson et al., 2019). Understanding whether EVs represent a dispensable byproduct or an essential effector of paracrine senescence could inform the design of more precise interventions. Moreover, given that EVs are membrane-bound and potentially druggable through distinct mechanisms compared to soluble factors, identifying their specific contributions may reveal new therapeutic vulnerabilities.

In this study, we systematically investigate the role of EVs secreted by senescent tumor cells in mediating paracrine senescence. Using bleomycin-induced senescent A549 lung adenocarcinoma cells as a model, we demonstrate that conditioned medium from senescent cells induces senescence markers in recipient tumor cells, including altered morphology, senescence-associated β-galactosidase (SA-β-GAL) activity, and upregulation of the cyclin-dependent kinase inhibitor *CDKN1A*. Through both pharmacological inhibition of EV biogenesis using GW4869 and genetic downregulation of the EV secretion mediator *RAB27A*, we establish that reduction of EV release markedly attenuates paracrine senescence without affecting soluble SASP factor secretion. Proteomic characterization of senescent EVs reveals distinct molecular signatures consistent with previously reported senescent vesicle profiles. Critically, EVs isolated by size-exclusion chromatography retain full paracrine senescence-inducing capacity in the absence of soluble SASP components. These findings establish senescent EVs as autonomous mediators of paracrine senescence and highlight their potential as specific targets for modulating the senescence secretory response in cancer and aging.

## RESULTS

### Conditioned media from senescent A549 cells induce paracrine senescence

Secreted factors from senescent primary cells, collectively termed senescence-associated secretory phenotype (SASP), have been previously reported to be capable of inducing senescence in neighboring cells (Acosta et al., 2013; Acosta et al., 2008; Kuilman et al., 2008). To test the activity of secreted factors from senescent tumor cells, we decided to analyze the conditioned media (CM) derived from senescent human lung adenocarcinoma A549 cells. For this, A549 cells were treated with bleomycin, a DNA damaging agent widely used to induce senescence (Aoshiba et al., 2013; Aoshiba et al., 2003). Treated cells exhibited changes in cell morphology, an increase in SA-β-GAL activity, a higher expression of *CDKN1A*, and decreased proliferation **(Fig. 1a and Supplementary Fig. 1a-f).** In addition, the transcriptomic analysis of these bleomycin-treated A549 cells revealed a clear separation of treated and untreated samples, with the treated cells clustering together and sharing a senescence profile. This was characterized by the altered expression of differentially expressed genes typically found during cell senescence, such as the increase in *CDKN1A*, *SERPINE1*, *MDM2*, or *FAS*, or the decrease in *FOXM1*, *CDK1*, *LMNB1* or ***PLK1*(**Fig. 1b-e and Supplementary Fig. 1g-j).

**Fig. 1.**
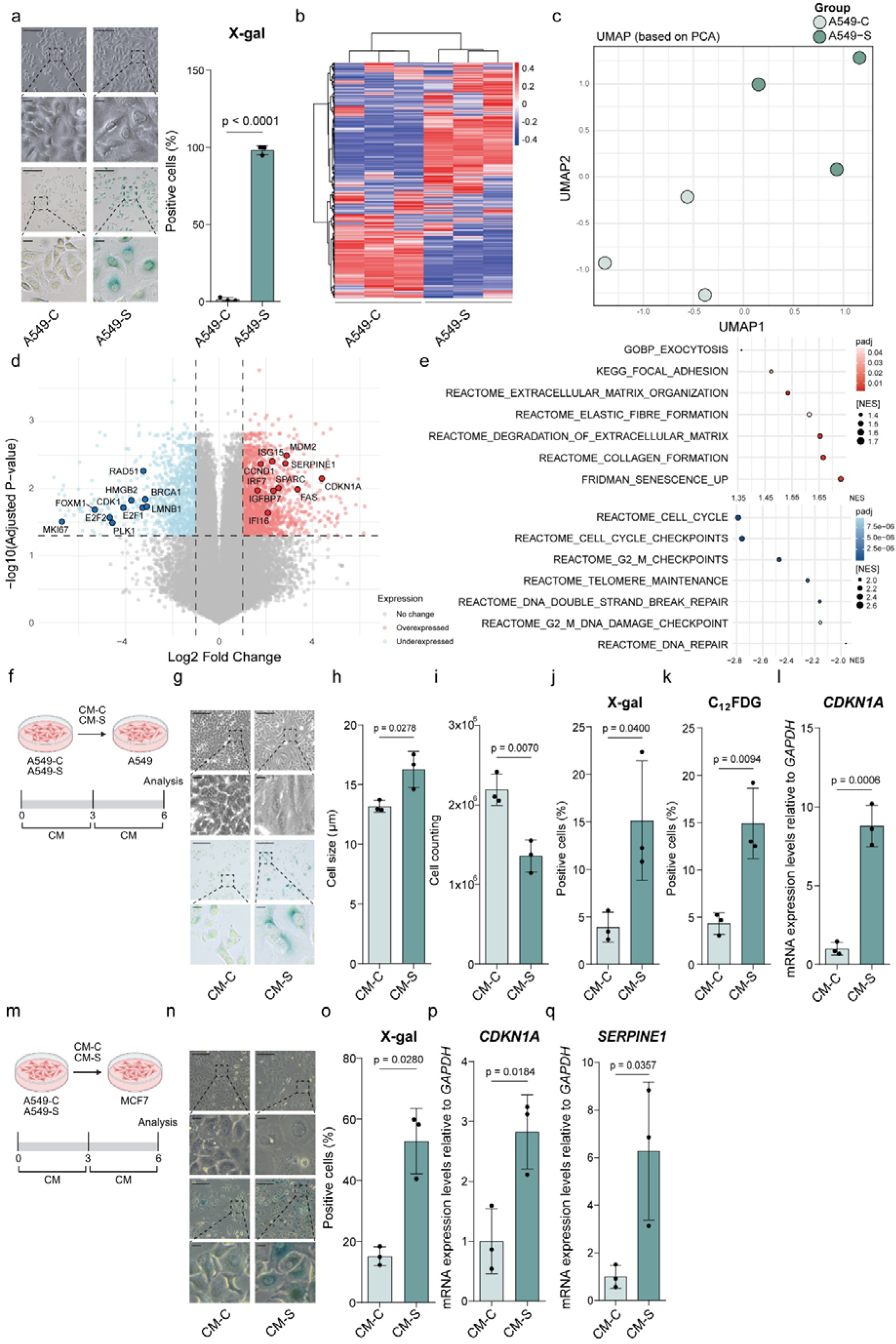
Conditioned media from chemotherapy-induced senescent A549 cells induce paracrine senescence in proliferative A549 cells. (a) Representative images of cell morphology and SA-β-Gal staining of proliferative (A549-C) and senescent (A549-S) A549 cells (scale, 100 μm; scale in zoomed area, 200 μm). Quantification of the staining. (b) Heatmap representation of RNA-seq analysis of A549-C and A549-S. (c) UMAP showing group clustering. (d) Volcano plot of differentially expressed genes. (e) Dot plot analysis of differentially expressed pathways. Red dots represent upregulated pathways, and blue dots represent downregulated pathways. (f) Schematic representation of proliferative (CM-C) and senescent (CM-S) A549 CM treatment over proliferative A549 cells. (g) Representative images of cell morphology and SA-β-Gal staining, large scale, 200 μm, short scale, 100 μm. (h) Cell size measurement. (i) Cell counting. (j) Quantification of SA-β-Gal positive cells using X-gal substrate. (k) Quantification of SA-β-Gal positive cells using C_12_FDG substrate. (l) RT-qPCR analysis of *CDKN1A*. *GAPDH* was used as housekeeping gene. (m) Schematic representation of proliferative (CM-C) and senescent (CM-S) A549 CM over proliferative MCF7. (n) Representative images of cell morphology and SA-β-Gal staining, large scale, 200 μm, short scale, 100 μm. (o) Quantification of SA-β-Gal positive cells using X-gal substrate. (p) RT-qPCR analysis of *CDKN1A*. *GAPDH* was used as housekeeping gene. (q) RT-qPCR analysis of *SERPINE1*. *GAPDH* was used as housekeeping gene. Significance was calculated using Student’s t-test showing the exact p-value. Less than 0.05 was considered statistically significant (n= 3).

We collected CM from control proliferative (CM-C) and senescent (CM-S) A549 cells and treated proliferative A549 cells as recipient cells. After two rounds of incubation with the CMs for three days each, we observed morphological changes in the cells receiving CM-S **(Fig. 1f)**. These cells exhibited an enlarged and flattened morphology and reached a lower confluency **(Fig. 1g-i)**. Since this was suggestive of an induction of cell senescence, we assessed these cell cultures for SA-β-GAL activity. Using both X-Gal and C_12_FDG as substrates, we observed a clear increase in SA-β-GAL positive cells after CM-S treatment **(Fig. 1j-k and Supplementary 1k).** Cells also presented an upregulation of *CDKN1A* expression **(Fig. 1l)**, a cell cycle inhibitor typically involved in the proliferative arrest that characterizes cell senescence. This observation was not restricted to A549 cells, but it was also observed when we used the human breast cancer cell line MCF7 **(Fig. 1m)**. Cells changed their morphology when they received CM-S from senescent A549 cells, were positive for SA-β-GAL activity and showed increased expression of *CDKN1A* and the SASP factor, *SERPINE1* **(Fig. 1n-q).**

Altogether, these results suggest that CM derived from senescent A549 tumor cells can induce paracrine senescence in neighboring tumor cells.

### Chemical inhibition of EVs biogenesis reduces paracrine senescence

The SASP is composed of a soluble fraction and extracellular vesicles (EVs). To study the relative contribution of these fractions to the induction of paracrine senescence, we used the chemical inhibitor of EVs biogenesis GW4869. This compound reduces the activity of N-SMase, an enzyme that catalyzes the synthesis of ceramide, a crucial component of EVs biogenesis (Catalano & O’Driscoll, 2020). First, we checked that the treatment with GW4869 did not alter the senescence phenotype of producer cells. Treated cells retained their morphological features and were positive for SA-β-GAL staining **(Fig. Supplementary 2a-d).** Then, we also wanted to confirm that GW4869 was only altering the production of EVs and was not affecting the production or release of soluble factors. For this, we incubated cytokine arrays with CM-C as a control, and senescent CM derived from senescent A549 cells (CM-S) or from senescent A549 cells treated with GW4869 to block EVs biogenesis (CM-S-GW4869). CM-S was enriched in several of the cytokines, chemokines and growth factors that have been reported to be part of the SASP compared to CM-C, as expected. When we incubated the arrays with CM-S-GW4869, we observed that this pattern of soluble factors was unaltered by the treatment with GW4869, indicating that it does not modulate the release of the soluble factors of the SASP **(Fig. 2a).** On the other hand, to check the inhibition of EVs biogenesis by the compound, we measured the number of vesicles in the different CMs by nanotracking analysis (NTA). When we analyzed CM-S, we observed a clear increase in the number of particles detected by NTA compared to CM-C. This is in agreement with several reports that found an increase in the number of EVs released by senescent cells. In contrast, when we analyzed CM-S-GW4869 we observed a drastic reduction in the number of EVs **(Fig. 2b).**

**Fig. 2.**
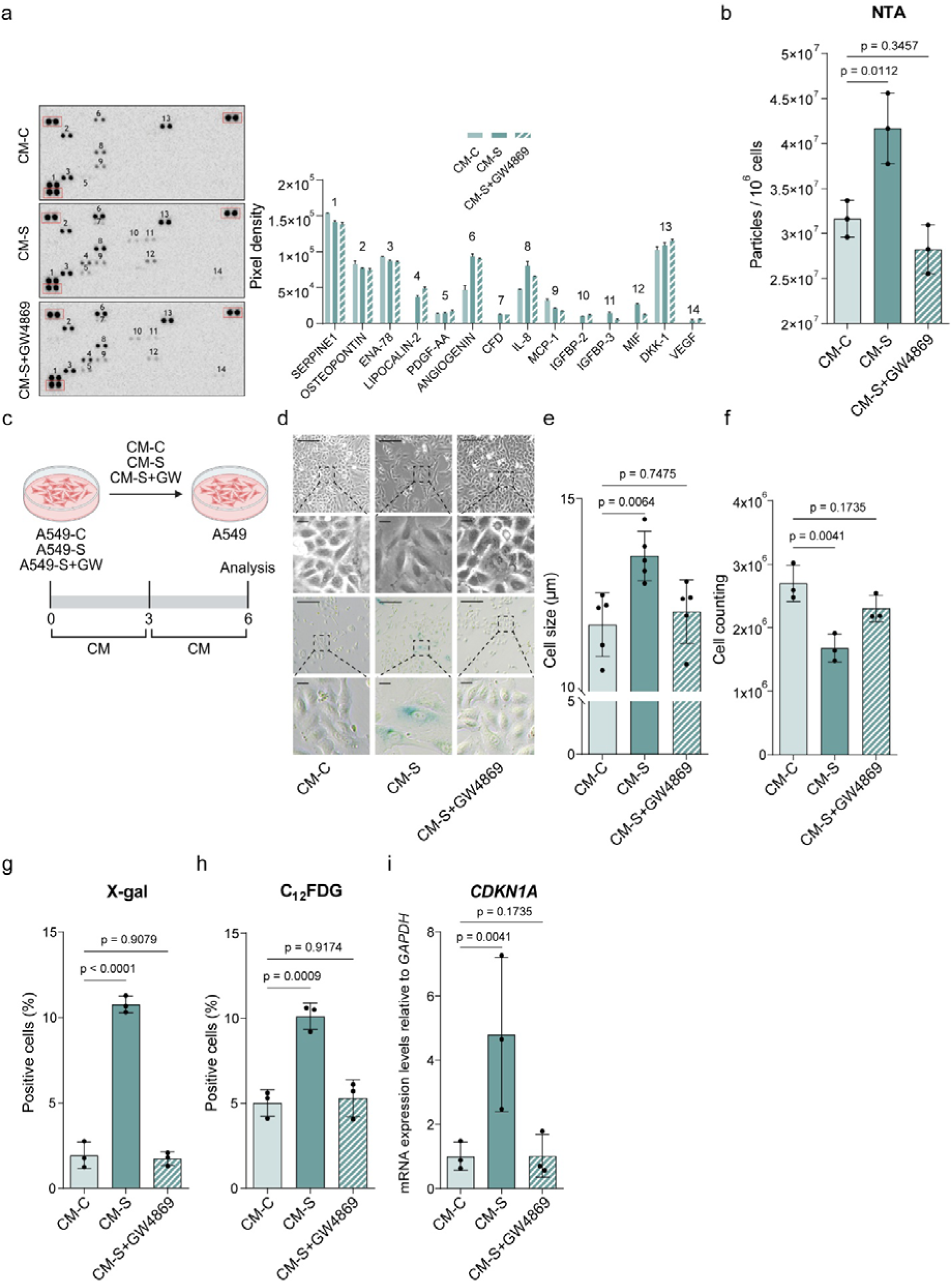
Chemical inhibition of EV biogenesis by GW4869 treatment reduces paracrine senescence. (a) Cytokine array analyses of CMs from proliferative (CM-C), senescent (CM-S), and GW4869-treated senescent cells (CM-S+GW6849) and their quantification. Red squares mark reference dots. (b) NTA analysis of particles in the CM. (c) Schematic representation of treatments using CMs on proliferative A549. (d) Representative images of morphology and SA-β-Gal staining (scale, 100 μm; scale in zoomed area, 200 μm). (e) Cell size measurement. (f) Cell counting. (g) Quantification of SA-β-Gal positive cells using X-gal substrate. (h) Quantification of SA-β-Gal positive cells using C_12_FDG substrate. (i) RT-qPCR analysis of *CDKN1A*. *GAPDH* was used as housekeeping gene. Significance was calculated using one-way ANOVA showing the exact p-value. Less than 0.05 was considered statistically significant (b-i, n=3; a, n=1).

To test the functional consequences of a reduction in the number of EVs in our senescent conditioned media, we treated A549 cells with the different CMs **(Fig. 2c)**. Again, CM-S induced a clear senescent morphology and caused a decreased cell proliferation indicative of paracrine senescence induction. Accordingly, cells stained positive for SA-β-GAL activity and showed increased *CDKN1A* expression. In contrast, when cells were treated with CM-S-GW4869, containing fewer EVs, cells were morphologically more similar to the ones treated with control CM-C, were mainly negative for SA-β-GAL activity, proliferated as the control cells, and showed lower expression levels of *CDKN1A*. **(Fig. 2d-I and Supplementary 2e).**

Overall, these data suggest that chemically inhibiting EVs biogenesis by GW4869 leads to a decrease in the number of EVs and an alleviation of paracrine senescence. This effect of GW4869 is not caused by an alteration of the senescent phenotype of the producer cells nor by an alteration of the secretion of soluble factors.

### Genetic downregulation of EVs secretion alleviates paracrine senescence

To extend our results obtained with the chemical inhibitor GW4869, we took advantage of the genetic downregulation of *RAB27A* expression to reduce the release of EVs. RAB27A mediates the fusion between the multivesicular body (MVB) and the plasma membrane, and its downregulation is associated with a reduction in the amount of secreted EVs (Ostrowski et al., 2010). To target RAB27A we used two shRNAs: shRAB27A #313 and shRAB27A #735. First, we characterized the reduction of *RAB27A* expression achieved by each construct through RT-qPCR and Western blot. While treatment with GW4869 had no effect on RAB27A levels, both shRNAs exhibited a robust capacity to reduce its expression, with shRAB27A #313 showing the strongest activity **(Fig. 3a-b)**. Importantly, neither #313 nor #735 shRNAs caused an alteration of the senescent phenotype of the cells as measured by SA-β-GAL staining using X-gal **(Supplementary 3c)**. To confirm the impact of *RAB27A* knockdown on the release of EVs, we measured the number of particles present in the different CMs by NTA **(Fig. 3c)**. While the CM from senescent cells expressing a non-targeting shRNA (CM-Scr-S) showed a higher number of particles compared to the control (CM-Scr), as expected, the senescent CM from cells expressing the two shRNAs targeting RAB27A (CM-shRAB27A #313-S and CM-shRAB27A #735-S) showed a clear reduction in the release of EVs.

**Fig. 3.**
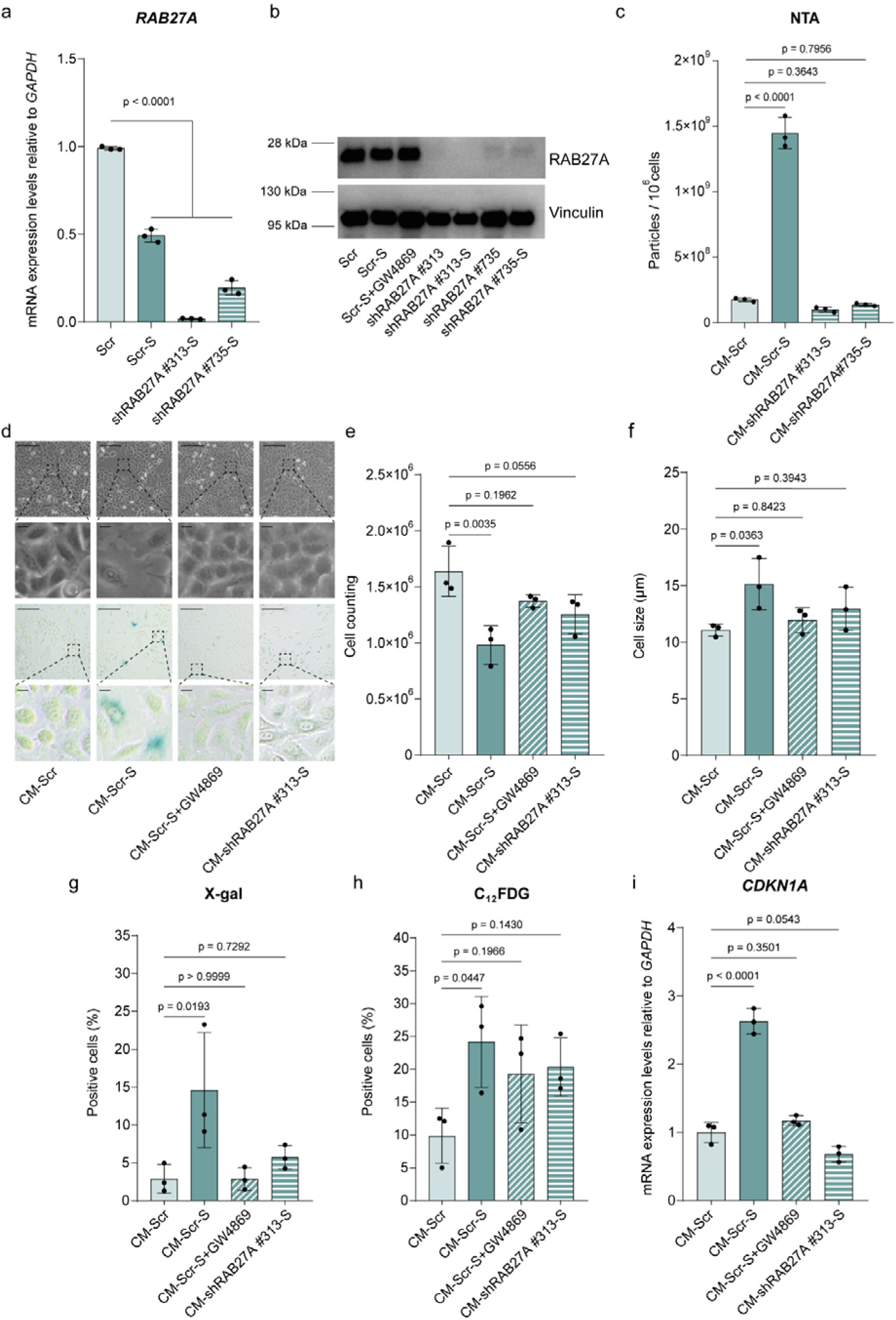
Genetic downregulation of RAB27A reduces EV secretion and paracrine senescence. (a) RT-qPCR analysis of *RAB27A*. *GAPDH* was used as housekeeping gene. (b) Western blot analysis of RAB27A. Vinculin was used as loading control. (c) NTA analysis of particles in CM. (d) Representative images of morphology and SA-β-Gal staining (scale, 100 μm; scale in zoomed area, 200 μm). (e) Cell counting. (f) Cell size measurement. (g) Quantification of SA-β-Gal positive cells using X-gal substrate. (h) Quantification of SA-β-Gal positive cells using C_12_FDG substrate. (i) RT-qPCR analysis of *CDKN1A*. *GAPDH* was used as housekeeping gene. Significance was calculated using one-way ANOVA showing the exact p-value. Less than 0.05 was considered statistically significant (a, c-i, n=3; b, n=1).

To analyze the effect of reducing the release of EVs by knocking down RAB27A, we added the CMs from control proliferating cells or after the expression of shRAB27A #313. Target cells incubated with CM-Scr-S reproduced the paracrine senescence induction already described, while both GW4869 treatment or RAB27A knockdown abolished this activity. Senescent cell morphology, reduction in cell numbers, SA-β-GAL activity, and an increase in *CDKN1A* expression only occurred when cells were incubated with CM-Scr-S but not with CM-Scr-S+GW4869 or CM-shRAB27A #313-S **(Fig. 3d-I and Supplementary 3c).**

Altogether, our data indicate that the reduction of EVs secretion by genetic downregulation of *RAB27A* alleviates paracrine senescence.

### Proteomic characterization reveals distinct features of senescent EVs

To obtain a deeper understanding of the senescent EVs, we decided to perform a proteomic analysis of EVs isolated from control proliferating or senescent A549 cells (C-EVs and S-EVs, respectively). Samples from both groups separated clearly on a UMAP **(Fig. 4a).** We managed to detect 1,844 proteins, 285 of them were significantly upregulated in S-EVs, and 267 were downregulated. We compared our proteomic profile with a previously described one obtained from EVs secreted by irradiation- or oncogene-induced senescence, and found several common upregulated and downregulated proteins, as highlighted in the volcano plot **(Fig. 4b)**. The differential protein expression analysis by cellular component revealed terms related with extracellular vesicles and the extracellular space, validating the identity of our purified EVs. When we analyzed gene ontology terms related with biological process, we found several activities previously reported for senescent EVs, such as “wound healing”, “blood coagulation”, or “hemostasis” **(Fig. 4c-e)**.

**Fig. 4.**
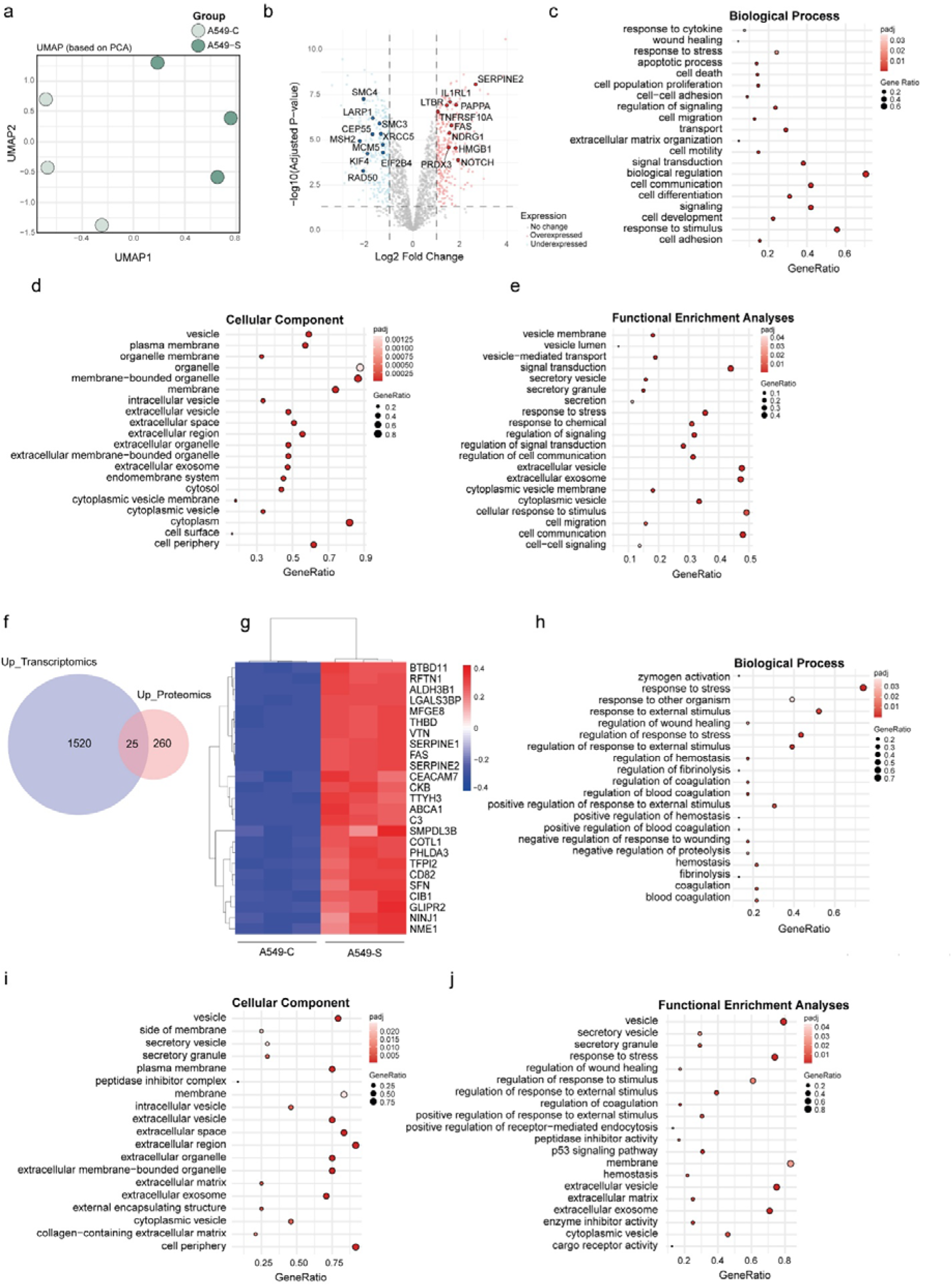
Senescent EVs exhibit a distinct proteomic profile. (a) UMAP showing group clustering of control (C-EVs) and senescent (S-EVs) EVs. (b) Volcano plot of differentially expressed proteins. (c) Dot plot analysis of differentially expressed Biological Process pathways, (d) Cellular Component terms, and (e) of Functional Enrichment Analyses pathways. (f) Venn’s diagram of upregulated transcripts (Up_transcriptomics) and proteins (Up_Proteomics) and their overlap. (g) Heatmap of overlapping proteins from Venn’s diagram. (h) Dot plot analysis of differentially expressed Biological Process terms of overlapping proteins, (i) Cellular Component pathways, and (j) of Functional Enrichment Analyses pathways.

Next, we decided to combine the transcriptomic and proteomic analyses. We identified a subset of 25 differentially upregulated proteins that were also transcriptionally increased **(Fig. 4f-g).** These 25 overlapping factors showed a strong upregulation at both the proteomic and transcriptomic levels, and were again related to extracellular vesicles and the extracellular space, as well as biological processes related to wound healing and hemostasis **(Fig. 4h-j).**

In summary, the EVs we isolated from bleomycin-induced senescent A549 cells exhibited proteomic features of previously described senescent EVs.

### Isolated senescent EVs can independently induce paracrine senescence

Our results showed that genetically or pharmacologically decreasing the number of EVs present in the CM from senescent cells abrogates the induction of paracrine senescence, suggesting that EVs may play an important role in this property of the SASP. To directly evaluate EV activity, we decided to isolate them from senescent CM by size-exclusion chromatography (SEC). Purified EVs (SEC-S-EVs) were analyzed by NTA to verify if their size corresponds to the expected range for EVs. Compared to particles from CM samples (CM-C, CM-S and CM-S-GW4869), SEC-S-EVs showed a similar median hydrodynamic diameter distribution (D50) and mode **(Fig. 5a).** Also, SEC-S-EVs were analyzed by transmission electron microscopy (TEM) and Western blot, showing that they express the canonical markers of EVs (TSG101, ALIX, Flotillin and CD81) and lack Calnexin, as expected **(Fig. 5b-c)**. To confirm that our EVs isolation method was efficiently removing soluble factors, we incubated cytokine arrays with our SEC-S-EVs preparation and compared the pattern of soluble factors with the one obtained by incubating with CM-S. We observed a drastic reduction in the number and intensity of soluble factors detected by this method, with most of them undetectable in the SEC-S-EVs condition **(Fig. 5d)**. These data suggest that our preparations contain purified EVs, dissected from the soluble factors of the SASP.

**Fig. 5.**
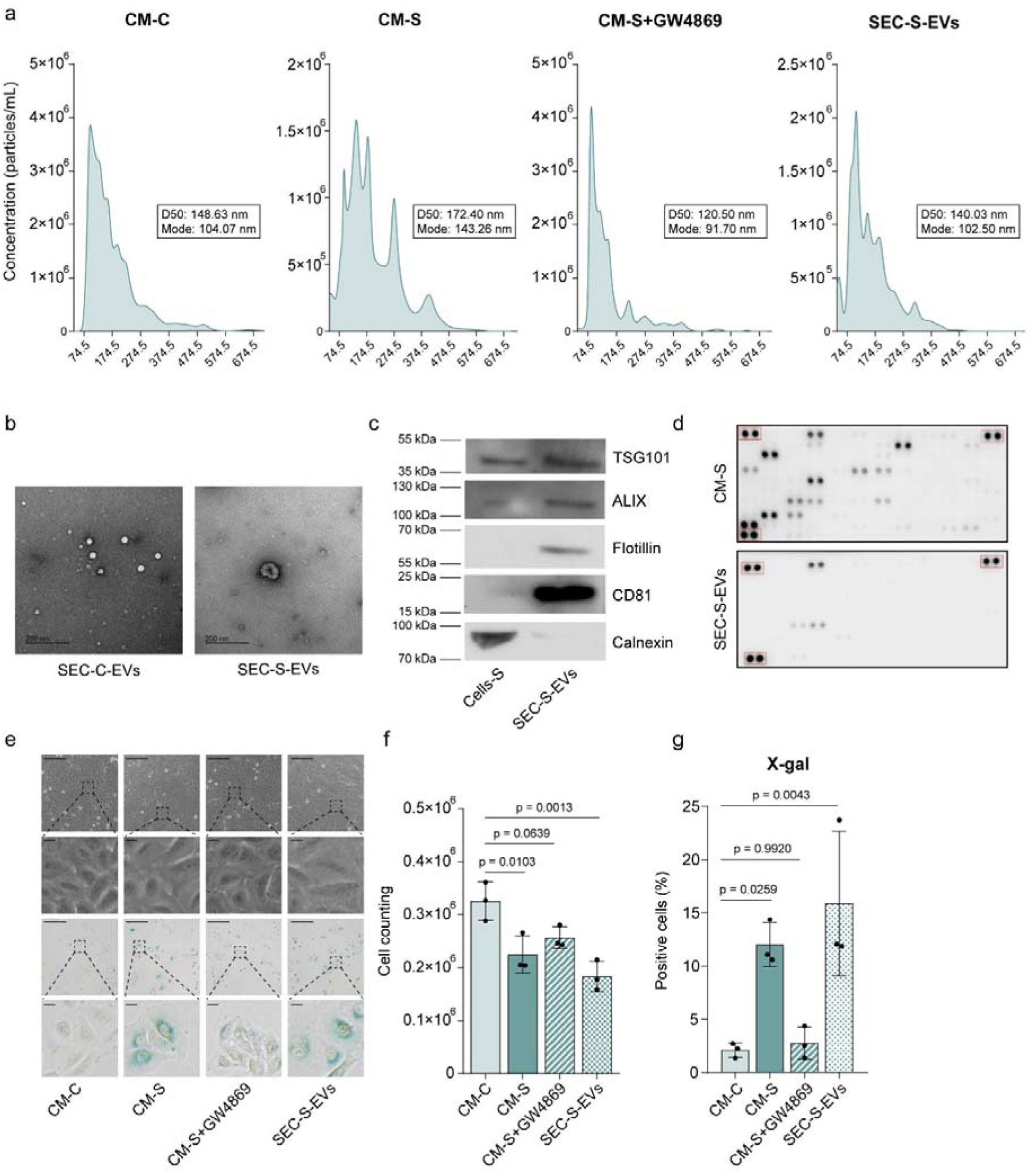
Isolated senescent EVs can independently induce paracrine senescence. (a) NTA analysis showing profiles, D50 and mode of particles within the CMs of control (CM-C), senescent (CM-S), senescent GW4869-treated (CM-S+GW4869) cells, and EVs isolated by SEC from senescent cells (SEC-S-EVs). (b) Transmission Electron Microscope (TEM) images of EVs isolated by SEC from control (SEC-C-EVs) and senescent (SEC-S-EVs) cells. (c) Western blot analysis showing the presence of canonical markers of EVs (TSG101, ALIX, Flotillin and CD81) and absence of ER marker calnexin in EVs isolated from senescent (SEC-S-EVs) cells. (d) Cytokine array analysis of soluble proteins in CM-S and SEC-S-EVs. (e) Representative images of morphology and SA-β-Gal staining (scale, 100 μm; scale in zoomed area, 200 μm). (f) Cell counting. (g) Quantification of SA-β-Gal positive cells using X-gal substrate. Significance was calculated using one-way ANOVA showing the exact p-value. Less than 0.05 was considered statistically significant (a, e-g, n=3; b-d, n=1).

To directly assess the potential activity of the EVs isolated from senescent cells, we added SEC-S-EVs to actively proliferating A549 cells and compared senescence induction with CM-S. The treatment with CM-S induced morphological changes resembling cell senescence and caused a reduction in cell proliferation as well as SA-β-GAL positive staining, while the pharmacological inhibition of EVs (CM-S-GW4869) reproducibly abrogated this effect. In contrast, the treatment with SEC-S-EVs recapitulated the same effects as the complete CM-S **(Fig 5e-g).**

Taken together, these results reinforce the observation that the EVs from senescent cells can induce paracrine senescence independently of the rest of the components of the SASP.

## DISCUSSION

The senescence-associated secretory phenotype is increasingly recognized as a multidimensional signaling system with soluble and vesicular arms that act in concert to remodel tissues and influence disease course (Basisty et al., 2020; Tanaka & Takahashi, 2021; Wang et al., 2024). By integrating pharmacologic inhibition, genetic perturbation, unbiased proteomics, and functional reconstitution, our study delineates a central, nonredundant role for extracellular vesicles in transmitting senescence among tumor cells. Specifically, we show that reducing EV production pharmacologically or genetically attenuates the paracrine senescence activity of bleomycin-induced senescent A549 cells without affecting the soluble SASP profile, while at the same time, purified senescent EVs are sufficient to drive paracrine senescence. Our data indicate that EVs are not merely correlates of the senescent state but active mediators of its propagation.

Prior work has established that soluble SASP components can induce senescence in neighboring cells (Acosta et al., 2013). However, the necessity and sufficiency of EVs have been less clear, especially in cancer contexts. Our experimental strategies converge on the same conclusion: depleting EVs curtails paracrine senescence, whereas purified senescent EVs restore it. The observation that GW4869 reduces EV number without altering producer-cell senescence or soluble SASP composition strengthens the causal link to vesicular cargo rather than confounding changes in the global secretome. Likewise, *RAB27A* knockdown, which diminishes EV release via impaired fusion of the multivesicular body with the plasma membrane, phenocopies the pharmacologic intervention, consolidating the mechanistic interpretation. Thus, our results show that EVs are necessary and sufficient mediators of paracrine senescence.

Proteomic characterization revealed that senescent EVs show a distinct proteomic profile. Senescent EVs harbor a signature enriched for extracellular vesicle/extracellular space components and biological processes such as wound healing, hemostasis, and tissue remodeling. This profile aligns with the broader concept that senescence can orchestrate microenvironmental changes conducive to both repair and pathology. The overlap between transcriptional upregulation and EV protein enrichment highlights candidate effectors whose biogenesis is coordinated at multiple regulatory layers. These factors may function as proximal drivers of the senescence response in recipient cells and serve as biomarkers of EV-mediated signaling.

Therapy-induced senescence (TIS) is a common outcome of genotoxic treatments and can paradoxically promote tumor relapse and resistance through persistent SASP signaling (Prasanna et al., 2021; Song et al., 2023; Wang et al., 2024). Our findings suggest that vesicular components of the SASP represent actionable levers to modulate TIS consequences. For instance, targeting EV biogenesis or release (e.g., via *RAB27A* pathways) could reduce the propagation of senescence to proliferating tumor cells and stromal elements, potentially curtailing the establishment of pro-tumorigenic niches. Conversely, in settings where senescence reinforcement is desired, for example, in combination with immune-mediated clearance, strategies to enhance or engineer EV cargo might be leveraged therapeutically.

In summary, our results define senescent EVs as necessary and sufficient effectors of paracrine senescence in tumor cells and unveil a distinct proteomic program associated with their activity. These insights refine the architecture of the SASP and provide a framework for targeting vesicular pathways to modulate senescence-driven phenotypes in cancer.

Together, these results uncover a central role for EVs in mediating senescence propagation among tumor cells and provide mechanistic insight into how senescent cells influence their microenvironment. This work advances the understanding of SASP heterogeneity and identifies senescent EVs as key regulators of non–cell-autonomous senescence responses with potential implications for cancer progression, therapeutic resistance, and senescence-targeting interventions.

## AUTHOR CONTRIBUTIONS

V.E.-S., A.M. and S.D.S.-A. performed most of the experimental work; P.P., P.L.-F., M.A.P., A.F.-F., R.P.-P., and M.G.-B. helped with experiments and reagents; J.R.-O., M.J.A., M.M., and E.P. helped with the analysis of EVs; G.N.C. and C.S.M. helped with TEM; P.X.-E. and J.M. performed proteomics; R.A.-V., E.V.D.L. and A.D. performed transcriptomics; V.E.-S., M.C., and S.D.S.-A. wrote the manuscript; M.C., and S.D.S.-A. conceptualized and supervised the project.

## ACKNOWLEDGEMENTS

Work in the laboratory of M.C. is funded by grants from MICIU/AEI/10.13039/501100011033 and by “ERDF/EU” (PID2021-125479OB-I00 and PID2024-159335OB-I00) and GAIN, Xunta de Galicia (IN607B2024/13). S.D.S.-A. is supported by a Juan de la Cierva fellowship (JDC2022-049032-I) funded by MCIU/AEI/10.13039/501100011033 and European Union “NextGenerationEU”/PRTR”. M.A.P. was supported by predoctoral fellowships from GAIN, Xunta de Galicia (ED481A 2022/432) and FPU program from MCIU (FPU23/02493). A.M. was supported by a predoctoral fellowship from FPU program from MCIU (FPU24/00846). R.P-P. received a fellowship from GAIN, Xunta de Galicia (ED481A-2025-101)

We acknowledge the support of the María de Maeztu program grant (CEX2024-001463-M) funded by MICIU/AEI/10.13039/501100011033, and from the “Regional Ministry of Education, Science, Universities and Vocational Training” of the Xunta de Galicia through the CIGUS Network of Research Centres (ED431G/2023/02), and the European Union through the European Regional Development Fund (ERDF).

## METHODS

### Cell line generation and reagents

A549 (RRID:CVCL_0023) and MCF7 (RRID:CVCL_0031) cell lines were obtained from the ATCC. Both cell lines were cultured in high-glucose DMEM (4500 mg/L, Sigma-Aldrich) supplemented with 10% FBS (Sigma-Aldrich), 1% penicillin-streptomycin (Sigma-Aldrich), 1% L-glutamine (Sigma-Aldrich), and maintained in an incubator at 37 °C and 5% CO_2_ atmosphere. Cells were routinely tested for mycoplasma. Non-targeting scramble plasmid (Scr), shRAB27A #313 and shRAB27A #735 were a kind gift from Dr. Alberto Fraile-Ramos (Fraile-Ramos, A. et al., 2010). For lentiviral transduction, 5×10^6^ HEK-293T cells were seeded 24 h before transfection with a plasmid mixture in the presence of polyethyleneimine reagent 1 mg/mL (PEI, Polysciences) in a 1:1 plasmid DNA: PEI proportion. Third-generation lentiviral packaging plasmids pLP1, pLP2 and pLP-VSVG (ViraPower Lentiviral Packaging Mix, Invitrogen) were used. 36 h post-transfection, we collected lentiviral-enriched conditioned media every 12 h to treat target A549 cells for three consecutive rounds. The conditioned media were filtered through a 0.45 μm filter and supplemented with 8 μg/mL of polybrene reagent (Sigma-Aldrich). We treat A549 cell line with these conditioned media and selected with puromycin (Millipore) at 1 μg/mL.

For chemotherapy-induced senescence we used bleomycin (Mylan pharmaceutics) at 20 μM for 5 days. After senescence induction, conditioned media were collected following 24 h of incubation with complete DMEM. Conditioned media were centrifuged for 10 minutes at 500 g to deplete cellular debris and apoptotic bodies and stored at 4 °C until use.

For GW4869 treatment, cells were treated at 10 µM for 24 h following senescence induction. Then, cells were washed with 1x PBS and media was conditioned for 24 h.

### RT-qPCR

For gene expression analysis, we extracted total RNA from cell cultures using the Nucleospin RNA Kit (Macherey-Nagel) following the manufacturer’s instructions. Then, we converted RNA into cDNA using High-Capacity cDNA Reverse Transcription Kit (Applied Biosystems). For each reaction, we used 33 ng of cDNA, oligonucleotides at a 0.25 μM final concentration, 5 μL NZYSpeedy qPCR Green Master Mix (2X), ROX (NZYTech), and nuclease-free water up to 10 μL total volume. Quantification was performed in the AriaMX Real-Time PCR System (Agilent Technologies). Primer sequences were:

*CDKN1A*: 5’-CCTGTAACTGTCTTGTACCCT-3’, 5’-GCGTTTGGAGTGGTAGAAATCT-3’

*SERPINE1*: 5’-CTCATCAGCCACTGGAAAGGCA-3’, 5’-GACTCGTGAAGTCAGCCTGAAAC-3’

*RAB27A*: 5’-GCTTTGGGAGACTCTGGTGTA-3’, 5’-TCAATGCCCACTGTTGTGATAAA-3’

*GAPDH*: 5’-TCCATGACAACTTTGGCATCGTGG-3’, 5’-GTTGCTGTTGAAGTCACAGGAGAC-3’

### EVs isolation

For EVs isolation, we followed the instructions of Smart SEC^TM^ Single for EV Isolation kit (System Biosciences). For functional analysis, we resuspended EVs preparations in DMEM supplemented with 40% FBS, 4% penicillin/streptomycin and 4% L-glutamine in a 3:1 proportion to reconstitute the standard complete media.

### NTA characterization

To determine particle concentration, conditioned media was analyzed by nanoparticle tracking analysis (NTA). For direct conditioned media analysis, samples were diluted 1:10 in 1x PBS, whereas isolated EVs were diluted 1:100 in 1x PBS. Measurement was performed using Nanosight NTA v2.3 (Malvern Instruments). We used a detection threshold of 11 and a camera level of 15.

### Cell morphology and proliferation analysis

For cell counting and size, cells were trypsinized, resuspended in DMEM supplemented with 10% FBS and analyzed by automatic cell counter Luna II (Logos Biosystems). Cell morphology images were taken in an Axio Vert.A1 Microscope (Zeiss).

### Cellular staining

For clonogenic assays, cells were seeded at low confluency and allowed to proliferate for 14 days. Then, cells were fixed with 4% paraformaldehyde (Electron Microscopy Sciences) and stained with 0.05% crystal violet solution (Sigma) for 30 minutes. After staining, cells were washed and crystal violet was eluted with 10% acetic acid. Optical density was measured at 570 nm using Epoch2 microplate reader (Biotek).

For SA-β-GAL activity detection, we used the chromogenic X-gal substrate. Cell culture was washed with 1x PBS and fixed with 2% paraformaldehyde (Electron Microscopy Sciences) and 0,2% glutaraldehyde for 15 minutes at room temperature. After fixation, cells were washed with 1x PBS and stained with X-gal solution (X-gal solution was prepared by mixing citric acid/sodium phosphate (pH 6.0) at 40 mM, NaCl at 150 mM, K₃Fe(CN)₆ at 5 mM, K₄Fe(CN)₆ at 5 mM, MgCl₂ at 2 mM, and X-gal (5-bromo-4-chloro-3-indolyl β-D-galactose) at 1 mg/mL in distilled water) at 37 °C overnight. Cells were washed at the end of the staining with 1x PBS. Images were taken using an Axio Vert.A1 Microscope (Zeiss). To quantify positive cells, 10 images per condition were analyzed.

For flow cytometry analysis of senescence, C_12_FDG substrate was used according to the instructions of the CellEvent™ Senescence Green Flow Cytometry Assay kit (Invitrogen). Samples were analyzed on a Cytoflex Flow Cytometer (Beckman Coulter), with 10,000 events recorded per sample and using FlowJo v10 software.

EdU incorporation assays were performed using Click-iT EdU Alexa Fluor 488 Imaging (Invitrogen) according to manufacturer’s instructions. Nuclei were counterstained with DAPI. A549 cells were incubated with 10 μM EdU for 2 h at 37 °C. Images were taken with Axio Vert.A1 Microscope (Zeiss) and 10 images per condition were quantified.

### Western blot

Cellular lysates were prepared in 1x RIPA buffer containing protease and phosphatase inhibitor cocktails (4693159001 and 4906845001, Sigma-Aldrich). To measure protein concentration, we used DC Protein Assay (BIO-RAD), following the manufactureŕs protocol.

A total of 25 μg of protein per sample was electrophoresed in 12% precast polyacrylamide gels (Invitrogen) using the XCell SureLock (Invitrogen) system and transferred to a 0.45 μm PVDF membrane (Merck-Millipore). Membranes were blocked with 5% milk solution and incubated with primary antibodies for GAPDH (sc-32233, Santa Cruz Biotechnology 1:1000), β-Actin (sc-8432, Santa Cruz Biotechnology, 1:1000), Vinculin (sc-76314, Santa Cruz Biotechnology 1:2000), RAB27A (#69295, Cell Signaling Technology, 1:1000), p53 (sc-6243, Santa Cruz Biotechnology, 1:1000), and p21 (#2947, Cell Signaling Technology, 1:1000), overnight at 4 °C. Then, membranes were incubated with HRP-conjugated secondary antibodies anti-mouse IgG-HRP (#31430, Invitrogen, 1:10000), anti-rabbit IgG-HRP (#31460, Invitrogen, 1:10000) for 1 h. Chemiluminescent detection was performed using the SuperSignal West Pico PLUS substrate (Thermo Scientific) in a ChemiDoc MP Imaging System (BIO-RAD).

To analyze EVs protein levels, EVs were lysed separately using 2% SDS. Protein concentration was determined using Micro BCA Protein Assay Kit (Thermo Scientific). A total of 25 μg of total protein was separated into 12% precast polyacrylamide gels for electrophoresis. After transferring the proteins into a PVDF membrane, the blot was blocked with 5% milk solution. Membranes were incubated overnight at 4 °C with the following primary antibodies: Alix (ab88743, Abcam, 1:500), Calnexin (Ab22595, Abcam, 1:1000), TSG101 (SC-7964, Santa Cruz Biotechnology, 1:1000), Flot-1 (SC74566, Santa Cruz Biotechnology, 1:100), and CD81 (SC166029, Santa Cruz Biotechnology, 1:500). Afterward, membranes were incubated for 1 h with HRP-conjugated secondary antibodies anti-mouse IgG-HRP (#31430, Invitrogen, 1:10000), anti-rabbit IgG-HRP (#31460, Invitrogen, 1:10000). Chemiluminescent detection was performed using the SuperSignal West Pico PLUS substrate (Thermo Scientific), and signals were visualized using radiographic X-ray films (FUJI).

### Cytokine array

Soluble factors released by senescent cells were analyzed by cytokine array following the instructions of the Proteome Profiler Human XL Cytokine Array Kit (Bio-Techne). Arrays were revealed using a ChemiDoc MP Imaging System (BIO-RAD).

### TEM

Copper grids coated with a collodion/carbon film were subjected to an electrical discharge for 15 s to enhance sample adhesion to the grid. Grids were incubated with the sample for 5 min at room temperature. Excess liquid was removed by blotting, and grids were washed with 50 µL of distilled water for 10 s. After a second blotting step, grids were stained with 2% uranyl acetate for 30 s.

### RNA seq library preparation, sequencing and generation of FastQ files

Total RNA (100 ng) was used to generate barcoded RNA-seq libraries using the NEBNext Ultra II Directional RNA Library preparation kit (New England Biolabs) according to manufacturer’s instructions. First, poly A+ RNA was purified using poly-T oligo-attached magnetic beads followed by fragmentation and first and second cDNA strand synthesis. Next, cDNA ends were repaired and adenylated. The NEBNext adaptor was then ligated followed by second strand removal, uracile excision from the adaptor and PCR amplification. The size of the libraries was checked using the Agilent 2100 Bioanalyzer and the concentration was determined using the Qubit® fluorometer (ThermoFisher)

Libraries were sequenced on a HiSeq4000 (Illumina) to generate 60 bases single reads.

FastQ files for each sample were obtained using bcl2fastq 2.20 Software software (Illumina).

### Proteomic analysis

EVs’ protein samples were obtained from FBS-free conditioned media. Samples were solubilized with Tris-HCl 100 mM, pH 8.0, 8 M urea solution and 10 μg of protein were digested with standard ASP1 protocol. Digested samples were processed by mass spectrometry (LC-MS/MS) through Ultimate 3000 RSLCnano (Dionex) system coupled with mass spectrometer Exactive HF-X (ThermoScientific). Data was processed with MaxQuantum (v 1.6.1.10) using standard configurations. We crossed our data with UniprotKB/Swiss-Prot, July 2018, 20373 sequences database. Label-free quantification was performed using the “match between runs” algorithm (match window of 0.7 minutes and alignment window of 20 minutes). The minimum peptide length was set to 7 amino acids, and a maximum of 2 missed tryptic cleavages was allowed. Results were filtered at a false discovery rate (FDR) of 1% at both the peptide and protein levels. Then proteinGroups.txt file was imported into Prostar (v1.14.7) for further statistical analysis. For the “Senescent versus Control” comparison, proteins with an absolute log2 fold change > 1 and an FDR-adjusted p-value < 0.05 (Limma) were considered differentially regulated.

### Exploratory Data Analysis (EDA) and functional enrichment tools

Raw normalized abundance values were imported from Excel files into R (v4.4.2). Missing values were imputed using a distribution-based approach prior to downstream analyses. Row-wise z-scores were calculated for visualization, and heatmaps were generated with hierarchical clustering (Euclidean distance, Ward’s method).

Principal Component Analysis (PCA) was performed on the variance-stabilized and transposed abundance matrix, and the top principal components were used to assess variation among experimental conditions. UMAP was applied to the leading principal components to visualize sample clustering in two dimensions.

Gene Set Enrichment Analysis (GSEA) was performed using ranked protein lists and curated gene sets from KEGG, Reactome, and senescence-related signatures. Significantly overexpressed proteins were further analyzed using the g:Profiler web tool for GO, Reactome, and KEGG enrichment. Results were visualized in R and overlap with the SASPAtlas database was assessed to identify known senescence-associated secretory phenotype (SASP) components.

### Statistical analysis

All data are represented as mean ± standard deviation. Statistical significance between two groups was calculated using Student’s t-test. For comparisons involving more than two groups, we used one-way ANOVA with Tukey’s post-hoc test. Statistical significance was represented as the exact p-value, considering less than 0.05 as statistically significant.

For differential expression analysis, log2 fold changes were calculated from group means and statistical significance was assessed using t-tests with Benjamini-Hochberg correction for multiple testing (FDR). Proteins with |log2FC| > 1 and adjusted p-value < 0.05 were considered significantly differentially expressed. Volcano plots were used for visualization.

Figure plots were generated with Graphpad Prism 9.4.1.

---

**Supplementary Fig. 1.**
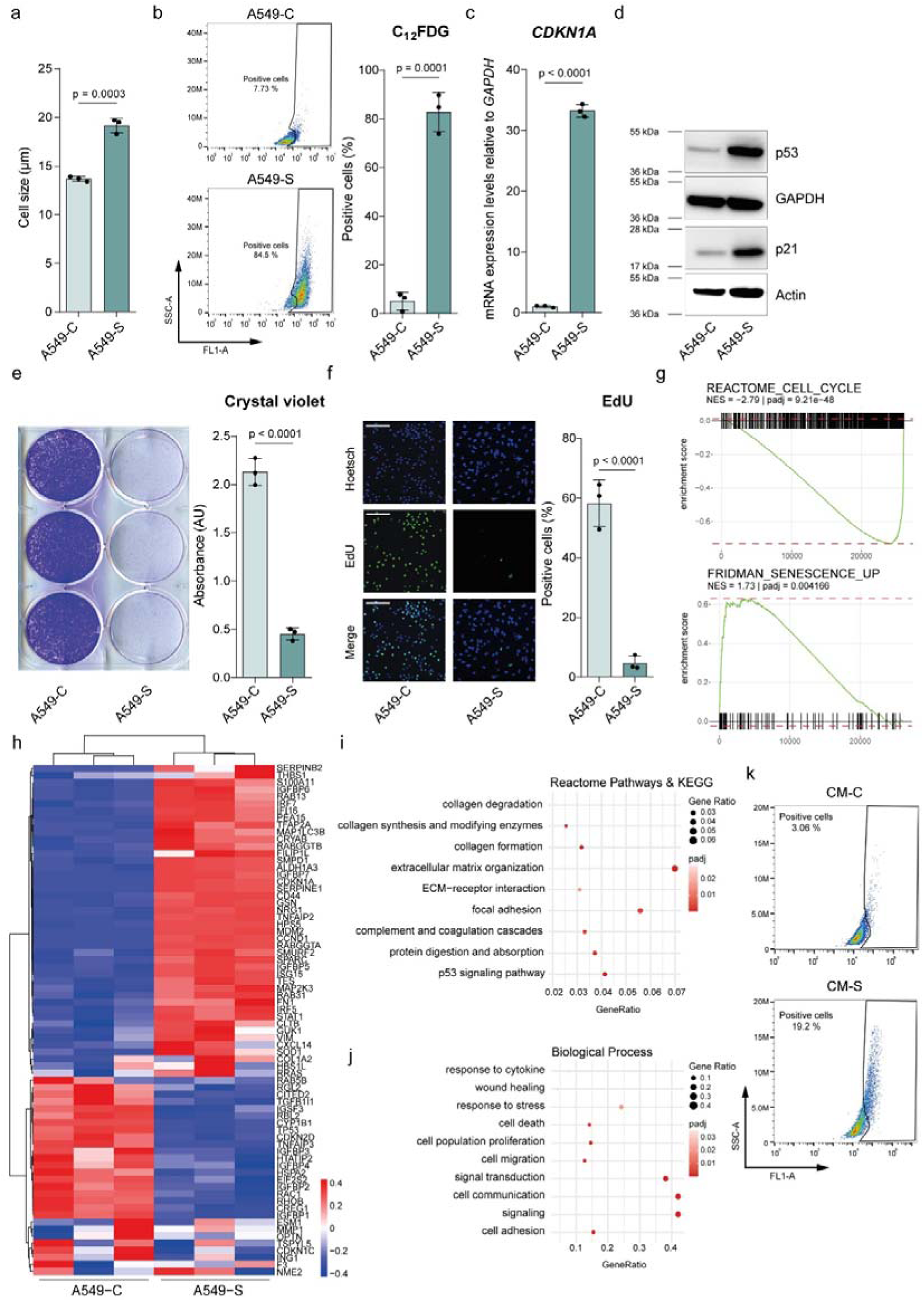
Characterization of chemotherapy-induced cellular senescence in A549 cells. (a) Cell size measurement of proliferative (A549-C) and senescent (A549-S) cells. (b) Flow cytometry plots and quantification of SA-β-Gal positive cells using C_12_FDG substrate. (c) RT-qPCR analysis of *CDKN1A*. *GAPDH* was used as housekeeping gene. (d) Western blot analysis of senescence markers (p53 and p21). GAPDH and Actin were used as loading controls. (e) Clonogenicity assay and quantification. (f) EdU incorporation assay and quantification (scale, 200 μm). (g) GSEA plots of the Reactome_Cell_Cycle and Fridman_Senescence_up gene sets. (h) Heatmap and list of differentially expressed genes. (i) Dot plot analysis of differentially expressed genes analyzed using the Reactome pathways and KEGG databases. (j) Dot plot analysis of differentially expressed genes using the Biological process terms in Gene Ontology. (k) Flow cytometry plots of cells treated with control (CM-C) and senescent (CM-S) conditioned media. Significance was calculated using Student’s t-test showing the exact p-value. Less than 0.05 was considered statistically significant (n=3, except d, n=1).

**Supplementary Fig. 2.**
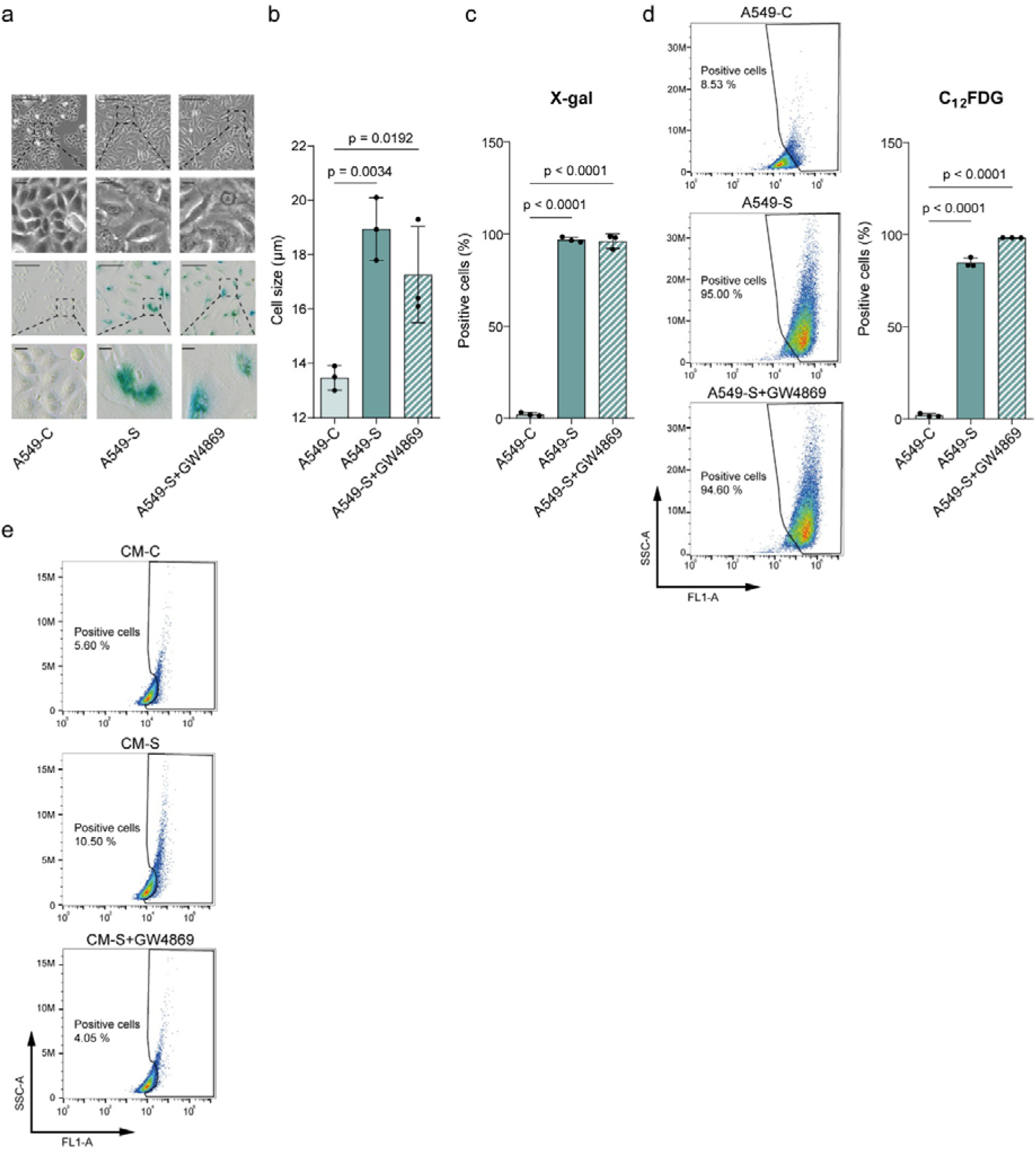
GW4869-treated senescent cells retain senescence phenotype. (a) Representative images of cell morphology and SA-β-Gal staining (scale, 100 μm; scale in zoomed area, 200 μm). (b) Cell size measurement. (c) Quantification of SA-β-Gal positive cells using X-gal substrate. (d) Flow cytometry plots and quantification of SA-β-Gal positive cells using C_12_FDG substrate. (e) Flow cytometry plots of cells treated with conditioned media derived from control (CM-C), senescent (CM-S) and GW4869-treated senescent cells (CM-S+GW4869). Significance was calculated using one-way ANOVA showing the exact p-value. Less than 0.05 was considered statistically significant (n=3).

**Supplementary Fig. 3.**
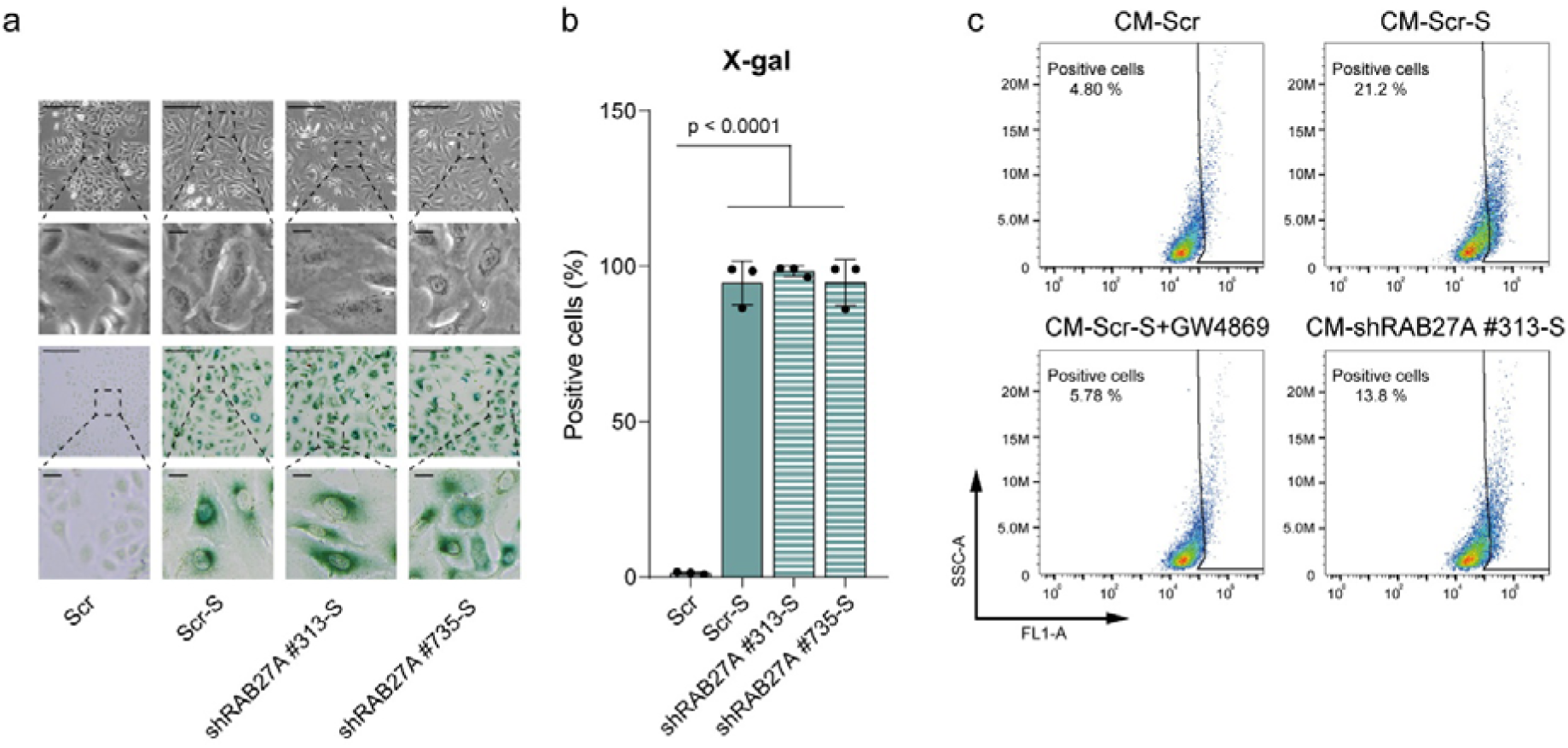
Senescence phenotype remains unaltered after RAB27A knockdown. (a) Representative images of morphology and SA-β-GAL staining (scale, 100 μm; scale in zoomed area, 200 μm). (b) Quantification of SA-β-GAL positive cells using X-gal substrate. (c) Flow cytometry plots of cells treated with conditioned media derived from control (CM-Scr), senescent (CM-Scr-S), or GW4869-treated senescent (CM-Scr-S+GW4869) cells, all of them expressing control Scrambled shRNA, and from RAB27A shRNA knockdown senescent cells (CM-shRAB27A #313-S). Significance was calculated using one-way ANOVA. Exact p-values are shown. Less than 0.05 was considered statistically significant (n=3).

